# Similar history biases for distinct prospective decisions of self-performance

**DOI:** 10.1101/607069

**Authors:** Ning Mei, Sean Rankine, Einar Olafsson, David Soto

**Author notes:** Correspondence to, Basque Center on Cognition, Brain and Language, Paseo Mikeletegi 69, 2nd Floor 20009 San Sebastian.

## Abstract

Metacognition can be deployed retrospectively -to reflect on the correctness of our behavior- or prospectively -to make predictions of success in one’s future behavior or make decisions about strategies to solve future problems. We investigated the factors that determine prospective decision making. Human participants performed a visual discrimination task followed by ratings of visibility and response confidence. Prior to each trial, participants made prospective judgments. In Experiment 1, they rated their belief of future success. In Experiment 2, they rated their decision to adopt a focused attention state. Prospective beliefs of success were associated with no performance changes while prospective decisions to engage attention were followed by better self-evaluation of the correctness of behavioral responses. Using standard machine learning classifiers we found that the current prospective decision could be predicted from information concerning task-correctness, stimulus visibility and response confidence from previous trials. In both Experiments, awareness and confidence were more diagnostic of the prospective decision than task correctness. Notably, classifiers trained with prospective beliefs of success in Experiment 1 predicted decisions to engage in Experiment 2 and vice-versa. These results indicate that the formation of these seemingly different prospective decisions share a common, dynamic representational structure.

## Introduction

The capacity to think about one’s own thoughts and behaviour is a fundamental constituent of the human mind which is known as metacognition. We rely on it to recognize that we have made a mistake, to realize that we have forgotten something important or to appreciate how confident we are about our own knowledge. An influential model of metacognition^1^ highlights the interplay between first-order, task-related processes (i.e., involving our perceptions and responses during task performance) and second-order processes, originating from the prefrontal cortex, that ‘monitor’ the correctness of the first-order process.^2, 3^

Monitoring of one’s own behavioural performance is typically assessed by means of retrospective reports, in which participants give ratings of confidence about their perceptual judgments or rate the state of visual awareness associated with the relevant stimulus. However, metacognitive processes are not just about thinking about one’s past and ongoing mental states. Metacognition can also be used prospectively to guide our future behaviour. For instance, we can mentally simulate ourselves in future probable scenarios, pre-empt the type of cognitive strategies needed to solve specific problems and adapt behaviour according to learning needs.

Prior research in the memory domain addressed how people make prospective judgments of learning during study,^4, 5^ and revealed, for instance, how decisions to study further rely on the evaluation of one’s own learning^6, 7^ and how this self-evaluation during study relates to subsequent memory accuracy. However, little is know about the factors that influence prospective metacognition during perceptual decision making. In addition to monitoring one’s own behavioral performance and forming prospective beliefs about future success, people also engage in self-regulation. For instance, people may also decide to put more attention when they lack confidence in their knowledge or stop further study when they are confident.

We here sought to investigate whether or not the formation of seemingly different types of prospective decisions (i.e. beliefs of success and decisions to engage with the environment) make use of similar information and recruit similar processes. Specifically, using a paradigm involving visual perceptual decisions we investigated the factors that predict future prospective beliefs of performance success vs. prospective decisions to engage with the environment (i.e. decisions to adopt a focussed attention state). Being successful in a higher proportion of recent trials may influence one’s estimation of prospective confidence, leading to predictions of a high probability of success in the next trial.^8^ A recent study by Fleming and colleagues (2016) investigated the formation of prospective and retrospective confidence.^9^ In this study, participants were asked to perform a motion discrimination task following by confidence ratings, and, every five trials participants also rated the prospective belief of success in the upcoming trial. The results showed that the prospective judgments were not associated with subsequent metacognitive performance (i.e. the association between confidence and accuracy). By contrast, in keeping with prior work, retrospective confidence judgments were closely aligned with task accuracy. Fleming and colleagues then modeled the influence of information in previous trials related to confidence and accuracy on the subjective estimates of prospective and retrospective confidence for a given trial. The results showed that current retrospective confidence can be predicted by the estimate of retrospective confidence in the previous trial. Prospective confidence was however dependent on the previous estimate of prospective confidence, and also on the prior retrospective confidence ratings over a longer timeframe (i.e. involving the previous four trials). The influence of task accuracy on prospective beliefs of success was far weaker by comparison.

Here we wondered whether different types of prospective decisions may rely on specific sources of information. After all, one’s certainty of the adequacy of one’s behavioural responses may be dependent on a host of different factors, including stimulus visibility, interference from distracting information and additional biases and heuristics.^10^ For instance, given a challenging perceptual task, a state of low visibility of the critical target may encourage the observer to decide to invest more effort in the next trial but may also lead to a reduction in confidence about his prospective accuracy. It is not known whether or not seemingly different types of prospective decision making (i.e. a decision to adopt a more focussed attention state on the next trial vs. prospective beliefs of success) are dependent on the same factors.

We modified an existing paradigm^11^ to quantify the contribution of the state of visual awareness, task-confidence and task-correctness to prospective decisions. We asked whether a similar or distinct pattern of experiences influenced different types of prospective beliefs, namely, predictions of success and decisions to engage with the environment. Participants were presented with an oriented Gabor patch near the threshold of visual awareness. Prior to the presentation of the Gabor, on each trial, participants indicated their belief of success (i.e. low or high) in the upcoming orientation discrimination task (Experiment 1) or indicated their decision to engage a focussed attention state (low or high; Experiment 2). Following the presentation of the Gabor, participants rated their visual awareness, responded to its orientation and rated their confidence in the orientation response.^11, 12^ Using standard machine learning algorithms, we sought to predict these seemingly different prospective decisions using information from previous trials concerning of awareness, task-confidence, and task-accuracy. We also evaluated the relative importance of these factors for prospective judgements using the coefficients from logistic regression. We also used a random forest classifier in order to estimate the stability of the decoding performance. Similar results were obtained. The results from the random forest classifier are presented in the Supplemental Materials.

Finally, we asked whether these seemingly different prospective decisions play a functional role in shaping our subsequent perception or metacognitive performance. We then tested whether the performance was affected by the type of prospective belief or decision to engage attention. The ‘self-fulfilling prophecy’ offers a view on the potential effect of prospective beliefs upon behavioural performance. Prospective beliefs may set an expectation that the participant is motivated to meet.^13, 14^ One possibility is that estimations of high probability of success may encourage observers to invest more cognitive resources in the upcoming trial and hence facilitate performance in a similar way to decisions to engage focussed attention. This study was devised to test these hypotheses.

## Experiment 1: Predictions of success

### Methods

#### Participants

Following informed consent, eighteen participants (19-23 years, mean age: 20.6, 6 males) took part in return of monetary compensation. This sample size was selected based on our prior study in which a similar paradigm was used.^11^ Data from three participants were excluded before analyses. One of the participants only reported a total of 3% of aware trials and two participants provided no responses in three conditions of awareness and confidence, impeding further analyses. This was likely due to inadequate pre-calibration of stimulus luminance (see below). The study conformed to the Declaration of Helsinki and was approved by the West London Research Local Research Ethics committee.

#### Experimental task and procedure

The experiment took place in a dimly lit room with a viewing distance of approximately 90 cm. The task was programmed and controlled by Psychopy.^15^ Stimuli were presented in a CRT monitor with a resolution of 1.600 x 1.200 pixels and a refresh rate of 85 Hz.

Figure 1 illustrates the experimental task. On each trial, participants were required to discriminate the orientation of a brief, masked Gabor, presented at the threshold of visual awareness. Prior to the presentation of the Gabor, participants reported their prospective belief of success associated with the upcoming trial (low vs high). Following the offset of the Gabor, participants rated their visual awareness of the Gabor, responded to its orientation and provided confidence ratings on the accuracy of the orientation responses.

**Figure 1:**
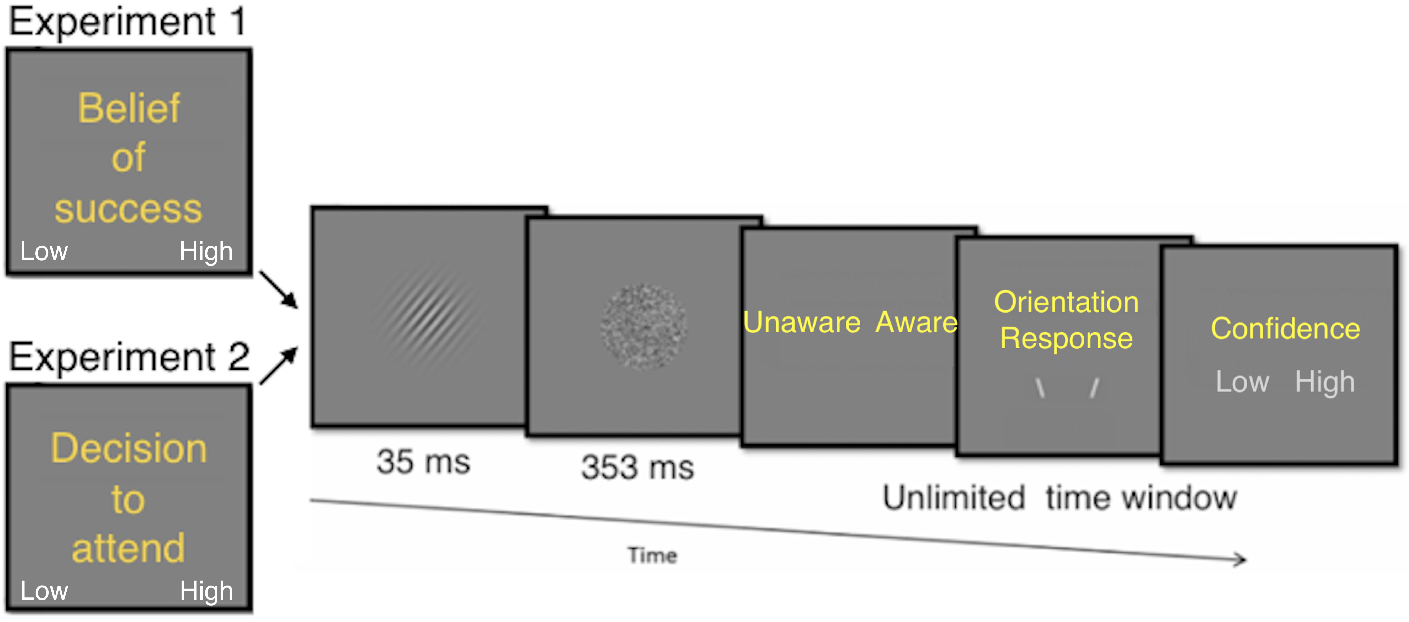
Illustration of the experimental protocol

During each trial, participants first made the prospective metacognitive decision during an unlimited time window. Then, a Gabor patch was presented in the center of the screen with a grey background (luminance = 10.48 cd/m^2^. Mask luminance was 11.34 cd/m^2^. The orientation of the Gabor was either 40 degrees to the left or right from vertical, and was randomly varied on each trial, with equal probability for each orientation. Participants responses were recorded using the keyboard.

Participants were instructed to complete a preliminary practice phase in order to get used to the orientation discrimination task. During this phase, the Gabor was presented for 362 ms with a fixed luminance of 11.93 cd/m^2^, and followed by immediate feedback regarding the accuracy of their response. Next, a calibration phase took place with a 35 ms Gabor stimulus duration and a 353 ms mask, similar to the experimental trials. Here, its luminance was varied using a staircase procedure. This meant that luminance increased when participants reported being unaware of the orientation of the Gabor and vice versa when they reported awareness. Participants were instructed to report ‘unawareness’ when they had no experience of the Gabor or saw a brief glimpse but otherwise had no awareness whatsoever of the orientation. They were instructed to report awareness of the Gabor when they could see its orientation somewhat or almost clear. The initial luminance was set to 11.97 cd/m^2^. The percentage of aware responses was computed on a trial to trial basis and the individual awareness threshold for each participant was determined by the luminance at the point where the probability of aware reports stabilized at around 0.5 for at least 10 trials, which were used to compute the average luminance to be used in the next procedural step.

Following the calibration, participants went through a training phase where they completed 15 trials identical to the experimental trials. The Gabor was presented for 35 ms during these trials. Prior to the presentation of the Gabor target, participants were instructed to report their belief of success in the upcoming task (high or low). Participants were not given specific instructions to report each belief equally often. Following the presentation of the target Gabor, participants reported their awareness of the Gabor, its orientation, and then rated their confidence in the orientation response. During the visibility response period, participants were presented with a screen displaying the response options (Unaware Aware). During the confidence response, period participants saw a screen with potential responses (Confidence: low high).

The justification for this particular order of responses is the following. Visual awareness of the stimulus was rated first to make sure that awareness was a genuine estimate of perceptual experience without being contaminated by memory interference from a longer delay between the stimulus and the awareness rating. The confidence judgment was given last because this referred to the orientation discrimination response, which followed the visual awareness response.

There was no response deadline for any of the three judgments. Participants were asked to provide precise ratings of awareness, and confidence, and accurate orientation discriminations without worrying about the speed of responding. Regarding the orientation response, participants were told that even if they were unaware of the stimulus, they should use their intuition and make their best guess about the orientation of the Gabor. Regarding the confidence report, participants were instructed to report how confident they were about the correctness of the orientation response on a relative scale of 1 (relatively less) to 2 (relatively more) confident. Participants were instructed that confidence ratings should be conceived in a relative fashion and hence they were asked to use all the confidence ratings independently of the awareness rating so that participants would not simply choose low confidence every time they were unaware and vice-versa on aware trials. Previous studies using a similar paradigm indicated that observers can display metacognitive sensitivity in both aware and unaware trials.^11, 12^

Prior to the experimental trials, there was a second calibration of the luminance of the Gabor starting with the luminance value from the first calibration. Each participant then completed 12 blocks of 50 trials (600 in total), with breaks between each block.

#### Machine learning protocols

We used standard machine learning algorithms (i.e. logistic regression and random forests models; Scikit-learn implementation) to predict whether the belief of success was low or high given a vector of features (i.e. correctness, visual awareness, and confidence) from the previous trials, considering 1-back, 2-back, 3-back, and 4-back trials, separately. Note that all the time series of trials back is not included in the classification. For instance, when we decode the belief of success based on the pattern of confidence, correctness and awareness of 4 trials back, we are only feeding the classifier with the data from that trial and do not include trials 1, 2 and 3 back. This range of trials was included based on a prior study,^9^ which showed that prospective confidence estimates were related to the confidence level four trials back, with this relatioship becoming stronger for trials closer to the current trial.

Scikit-learn by default implements the logistic regression with an L2 regularization built-in, which reduces the interpretability of the weight coefficients estimated by the regression. Therefore, we set the regularization to be very small in order to emulate no regularization as in the simplest form of logistic regression. During training, the model optimizer (LIBLINEAR), achieved the minimization of the difference between the predictions of the model and the true values we wanted to predict using a coordinate descent algorithm.^16^ Note the order of regressors does not matter because the Scikit-learn implemented logistic regression computes predictions by searching for the best-fit weights of the regressors using liblinear gradient descent algorithm.^16^ Since the weights, as parameters, are specific to the regressors, the order does not matter.

##### Cross-validation

We conducted a 100-fold shuffle splitting cross-validation for each subject, each decoding goal (1-back, 2-back, 3-back, and 4-back). Each fold was constructed by shuffling the examples. 80% of the data were selected to form a training set while the remaining 20% became the testing set. The predictive performance was estimated in each fold by comparing the target vector with the probabilistic predictions. The comparison was measured by the area under the receiver operating curve (ROC AUC). ROC AUC is a sensitive, nonparametric criterion-free, and less biased measure of predictive performance in binary classification,^17^ with 0.5 being the theoretical chance level. The AUC-ROC represents the ratio of the true positive classification rate (TPR, i.e. the classifier predicts ‘animal’ given the example is an animal) against the false positive rate (FPR, i.e. the classifier predicts ‘animal’ given the example is a tool).

##### Post-decoding

In order to estimate the significance level of the decoding performance, we generated a null distribution of decoding scores for each subject using a permutation analysis with 100 iterations. The null distributions were used as the empirical chance level. The null distribution of each subject was estimated by conducting the same cross-validation procedures with the same feature matrices and target vectors, except that the order of rows of the feature matrices and the target vectors were randomized independently. The average performance score over the permutation iterations represented the chance level estimate. This was found to be centered on the theoretical chance level of 0.5.

The statistical significance of the classification scores in each condition (i.e. trial back) was determined by using a non-parametric t-test. In it, the decoding scores were assessed relative to their corresponding chance level estimates. A permutation t-test was conducted to compute the uncorrected p-value for each trial back (1, 2, 3, 4), across all the subjects. However, for each experiment raw p values were corrected using Bonferroni multiple comparison correction procedure and compared to the nominal significance level of 0.05. The same applied for post-hoc tests after an ANOVA.

In plotting the classification results, the error bars represent bootstrapped 95% confidence intervals resampled from the average decoding scores of individual participants by the classifier with 1000 iterations.^18^

### Results

#### Classification analyses

First, we report the pairwise Phi correlations between the features used for classification. We note that since the correlation coefficient is inherently restricted to a range from −1 to +1, an arc-hyperbolic tangent transform^19^ was applied prior to statistical testing. The pairwise correlations were as follow: for (i) awareness and confidence: 0.224726 +/- 0.20307, t(14) = 4.28, p = 0.00226; (ii) correctness and awareness: 0.22878 +/- 0.10525, t(14) = 7.61, p = 0.000007; correctness and confidence: 0.16983 +/- 0.09330, t(14) = 6.61, p = 0.0000353. Although the correlations are higher than chance, they are far from 1, which is the maximal theoretical value that Phi can take. However, the maximal empirical value that Phi can take is likely to be less than 1 (i.e. the empirical distributions of the vectors for the two variables are unlike to be identical). In any case. these results indicate there is room for each variable to provide relevant information to classify the prospective belief of success.

We used a standard logistic regression classifier to predict the prospective belief of success on a given trial by using awareness, confidence, and correctness as training features from the preceding trials. We note that the probability of high beliefs of success was 0.5069 +/- 0.1507, which is not different from 0.5 (p = 0.57) and hence showing the overall the likelihood of each type of belief was similar. The results show that prospective belief of success could be predicted above chance levels using information from the previous trials. As shown in Figure 2, the prediction of success could be classified with the highest accuracy by using the features from the previous trial, with prediction accuracy dropping close to chance level based on information from 4 trials back. P-values following the permutation tests for the different classification analyses were as follows: 1-back: p < 0.0004, 2-back: p < 0.0004, 3-back: p < 0.0056, 4-back: p < 0.0084. Unless otherwise noted all p values reported are Bonferroni corrected for multiple comparisons. This pattern of results was also observed with a Random Forest Classifier (see Supplemental Figure 1).

**Figure 2:**
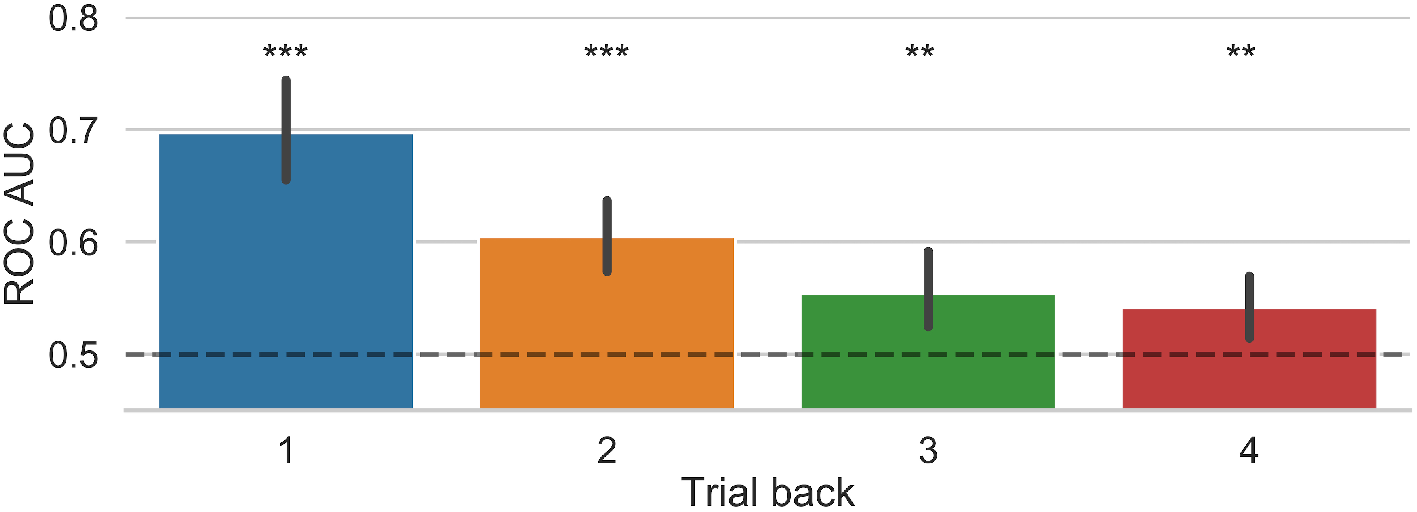
Results from the logistic regression classification model tested separately for each of trials back. Error bars represent bootstrapped 95% confidence intervals. ** p < 0.01, *** p < 0.001.

Next, we assessed the relevance of each of the different attributes (awareness, confidence, and correctness) for the classification. As we were interested in understanding the factors that contribute to future beliefs of success, we analyzed the weight coefficients (odd ratios) from the logistic regression (see Methods) by means of an ANOVA with time window (1,2, 3 and 4 trials back) and feature attribute as factors. Note that our main interest here is to understand the contribution of the different attributes for the classification. Since the above classification results already showed that classification accuracy decreases with the number of trials back, additional analyses based on significant main effects of time window are not considered further.

The analysis of the odd ratios of the logistic regression showed a main effect of window, F(3,42) = 19.15, p < 0.00001, *η*^2^ = 0.176, and a main effect of attributes, F(2,28) = 14.43, p = 0.00005, *η*^2^ = 0.179. Further t-tests showed that the odd ratios of both confidence (p = 0.0003449) and awareness (p = 0.0008536) was different from correctness. These results show that when observers rated high confidence/high awareness on the previous trial, then a belief of high success on the next trial was over 3 times more likely. Figure 3 illustrates this pattern of results. There was also an interaction between factors, F(6,84) = 10.66, p < 0.00001, *η*^2^ = 0.083. In the case of one trial back (N = 1), the odd ratio for confidence (p = 0.00012) and awareness (p = 0.0265) was higher than the odd ratio for correctness, but there was not difference between confidence and awareness (p = 1). In N = 2, the odd ratios of confidence and awareness were different from correctness (p = 0.00876 and p = 0.03797), but there was not difference between confidence and awareness (p = 1). This pattern of results was not observed for N = 3 and N = 4 (all ps > 0.08). Similar results were observed in the analyses of the feature importance of the Random Forest classifier (see Supplemental Figure 2). We also note that the variance of the odd ratios for the previous trial was larger than for increasing number of trials back. This is likely due to the odd ratios being closer to a value of 1 with increased numbers of trials back, meaning that features bear no influence on the prediction. The odd ratios displayed an effect of awareness and confidence at N-1 but the effect size is small and accordingly with a larger variance.

**Figure 3:**
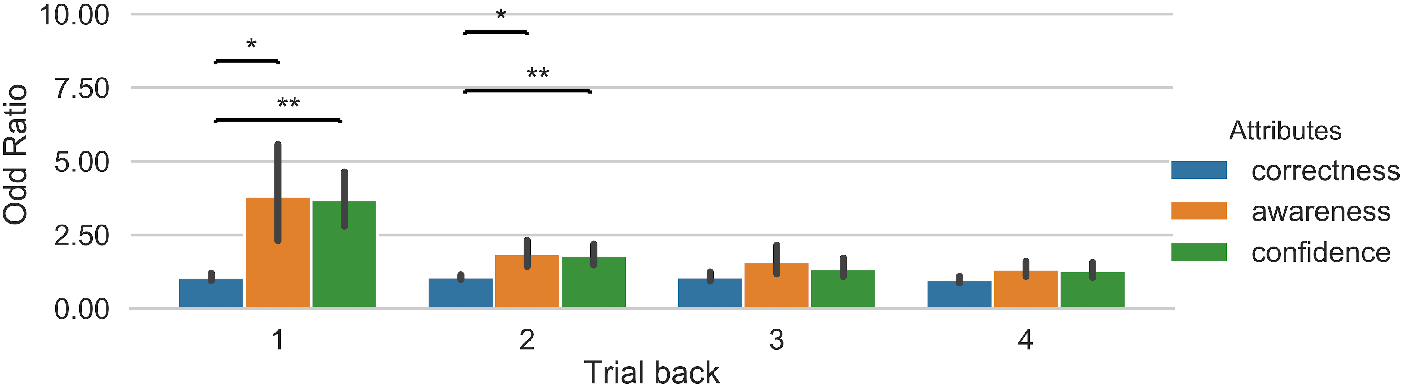
Illustration of odd ratios from the logistic regression for the different features involved in the classification of the belief of success. Odd ratios were computed separately for each of trials back. A value of 1 indicates that the odds of predicting the class is the same regardless of the feature value. Error bars represent bootstrapped 95% confidence intervals. * p < 0.05, ** p < 0.01.

#### Univariate analyses

We also performed univariate analyses of the probability of high success as a function of the experimental features of the previous trial. We used 1 trial back for the univariate analyses because here the multivariate classification results were strongest and thus provided an opportunity to verify the commonalities between the approaches. Accordingly, we calculated the conditional probability of high belief of success in the current trial as function of each level of correctness, awareness and confidence on the previous trial. Then we conducted a 2 (Correctness: Hit, Error) x 2 (Awareness: Aware, Unaware x 2 Confidence (high, low) repeated measures ANOVA on the probability of a belief of high success. There was no main effect of correctness on the probability of a belief of high success (F(1,13) = 1.703, p = 0.214). There was a main effect of awareness (F(1,13) = 14.091, p = 0.002) and also a main effect of confidence (F(1,13) = 62.429, p<0.001. The probability of a belief of a high success in the next trial was higher when the participants were aware of the stimulus on the previous trial and also when they were more confident. There was an interaction between correctness and awareness (F(1,13) = 5.796, p = 0.032). No other interactions were significant: F(1,13) = 0.535, p = 0.478, for correctness and confidence; F(1,13) = 0.924, p = 0.354, for awareness and confidence; and F(1,13) = 0.726, p = 0.409, for the three-way interaction. These results are in keeping with the classification analyses reported above. These results are shown in Supplementary Figure 3.

Further, we performed a linear mixed regression analysis using the lmerTest package^20^ to predict the prospective belief of success with fixed effects for each of the confidence, awareness and accuracy attributes considering the recent trial history up to 4 trials back with lagged factors (12 regressors in total) and random intercepts for each participant. The results indicate that confidence and awareness at 1-trial and 2-trials back predict the current prospective choice, which is also predicted from the confidence ratings at 3- and 4-trials back (see Supplemental Figure 4). This pattern of results is consistent with the multivariate classification approach, indicating that there there is information up to 4-trials back that is associated with the current prospective belief of success. Note that because the classification analyses were conducted for each trial back separately, it was unclear whether the prediction score based on information from several (e.g. 3) trials back was over-and-above that from 1 or 2 trials back. The linear mixed regression model including all the regressors addresses this issue (see also Fleming and colleagues^9^). Importantly, both univariate and multivariate classification approaches are complementary. For instance, in contrast with the univariate analyses in which all data is fit at once, the multivariate classification estimated model performance using cross-validation, hence allowing is to build a predictive model and assess its generalization to new samples. This is a clear advantage of the classification approach which will probe more critical for assessing the generalization of history biases across different prospective decisions in Experiment 1 (prospective beliefs of success) and Experiment 2 (decisions to engage attention), namely, whether the prior history of confidence, awareness and correctness in Experiment 1 predicts a seemingly different prospective choice in Experiment 2 (and vice versa). Assessing this cross-domain generalization is impossible to achieve using a standard univariate regression model in which all data is fitted at once.

#### Signal detection analyses

We computed type-1 d’ to index the observer’s sensitivity to discriminate the orientation of the Gabor, and also type-2 d’ or meta-d’ as a measu re of metacognitive sensitivity,^21^ which is basically a parametric estimation of the type-2 sensitivity (i.e. the capacity to discriminate correct from incorrect type-1 responses based on the confidence ratings) which is achieved by fitting a type-1 signal detection model to the observed type-2 performance and estimating the type-2 receiver operating characteristic (ROC) curves. In the type-1 model, the ‘signal’ and ‘noise’ were defined as left and right oriented Gabor, respectively. A ‘hit’ was, therefore, a correct response (‘left’) to a left-oriented Gabor and a ‘correct rejection’ was a correct response (‘right’) to a right-oriented Gabor. A ‘false alarm’ was an incorrect response (left) to a right-oriented Gabor and a ‘miss’ was an incorrect response (right) to a left-oriented Gabor.

A 2 x 2 ANOVA with awareness state (unaware, aware) and belief of success (high, low) as factors was carried out on the perceptual sensitivity to the orientation of the Gabor. There was an effect of awareness on type-1 d’ (F(1,14) = 52.44, p < 0.000004, *η*^2^ = 0.353). Perceptual sensitivity was higher on aware relative to unaware trials. There was no effect of prediction of success on perceptual sensitivity (F(1,14) = 0.021, p = 0.888, *η*^2^ = 0.00004) and no interaction effect between factors (F(1,14) = 0.096, 0.761, *η*^2^ = 0.0002). These results are depicted in Figure 4.

**Figure 4:**
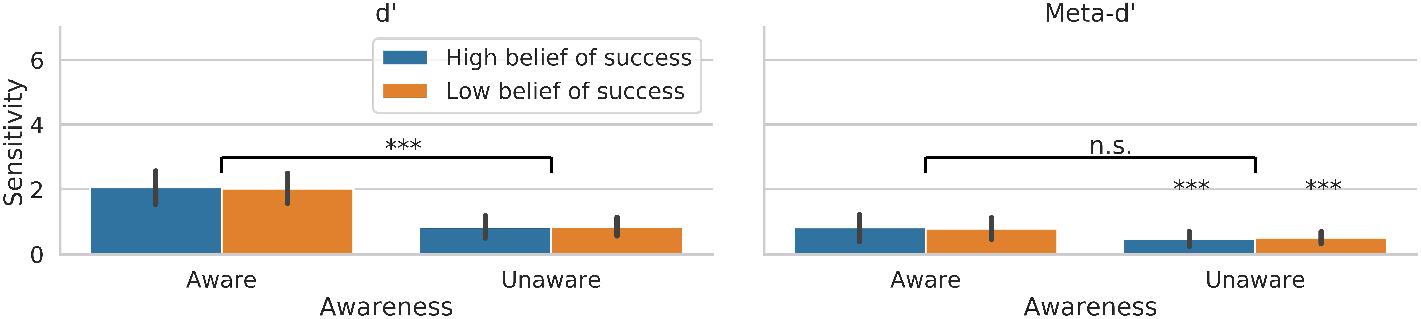
Perceptual sensitivity and metacogntive sensitivity scores as function of the state of awareness and the belief of success. Error bars represent standard error of the mean. n.s (not significant); *** p < 0.001.

A similar ANOVA was carried out on metacognitive sensitivity scores. There was no effect of awareness on type-2 d’ (F(1,14) = 3.273, p < 0.0919, *η*^2^ = 0.063), no effect of the belief of success (F(1,14) = 0, p = 0.988, *η*^2^ = 0.00002) and no interaction effect between belief of success and awareness (F(1,14) = 0.042, 0.84, *η*^2^ = 0.0008). Metacognitive sensitivity was well above chance even in trials that participants reported being unaware of the target (t(14) = 3.891, p < 0.0016 and t(14) = 5.218, p < 0.00013, for high and low belief of success cases).

There were no effects of the belief of success on M-ratio (meta-d’/d’), which is an index of metacognitive efficiency that factors out the level of d’^21^(F(1,14) = 2.78, p = 0.12. There was also no effect of awareness (F(1,14) = 0.007, p = 0.93) and no interaction between belief of success and awareness (F(1,14) = 2.44, p = 0.14).

### Discussion

We found that visual awareness had a profound effect on perceptual sensitivity, however, the effect of awareness on metacognitive sensitivity was far weaker. However, metacognitive sensitivity was well above chance across (un)awareness states. This replicates our prior observation^11^ and suggests that metacognitive operations are not necessarily carried out on the same type of representation or follow a similar process to that underlying first-order performance. In other words, the precision of retrospective metacognitive confidence judgments can be dissociable from factors that influence task performance (i.e. stimulus visibility). Hence, the results are in keeping with the proposal that metacognitive confidence and visual awareness can be dissociated (see also^22^) and with the view that higher-order cognitive processes are to some extent dissociable from conscious experience.^11, 12, 23–25^ However, further studies are needed in which metacognitive processing of perceptual decision making is assessed using both neural and behavioural measures under experimental conditions associated with null perceptual sensitivity.^26–28^

Most important for the aims of the present study, we observed that the current prospective belief of performance success could be predicted based on the pattern of visual awareness, task-confidence, and task-correctness, notably from several trials back. The prediction of this belief was strongest considering information from the most recent previous trial, with classification performance decreasing with longer time windows. Furthermore, we found that both confidence and awareness states, relative to task correctness, are more relevant for the classification of the prospective belief. However, this type of prospective beliefs did not appear to play a functional role in behavioural performance. Both perceptual sensitivity and metacognitive sensitivity were similar regardless of whether the prospective belief of success was high or low. This finding seems at odds with the possibility that prospective beliefs are associated with cognitive processing changes deployed to meet the belief (i.e. the ‘self-fulfilling prophecy’).^13, 14^ This raises the question of whether a different type of prospective decision (e.g. deciding whether to invest more attention on the current trial) may be predicted by the same information pattern that predicts prospective beliefs of success, and whether this type of prospective decision to engage with the environment may play a functional role in behavioural performance. Experiment 2 was designed with this in mind.

## Experiment 2: Decisions to engage attention

Experiment 2 was similar to Experiment 1 except that here a new set of participants reported their decision to engage in a more or less focussed attentional state on the upcoming trial, instead of making a prospective belief of their performance success. The goal of Experiment 2 was to test whether or not the findings of Experiment 1 generalize to this new context.

### Methods

#### Participants

Following informed consent, nineteen healthy volunteers (10 males and 9 females) took part in the experiment in return for monetary compensation. None of the participants took part in Experiment 1. They were aged between 18 and 47 years (mean age 21.6). Three participants were excluded prior to data analyses due to the absence of aware trials, likely due to inadequate stimulus calibration. All participants were right-handed and had normal or corrected-to-normal vision. They were naive to the experimental hypotheses and did not take part in Experiment 1. The study conformed to the Declaration of Helsinki and was approved by the West London Research Local Research Ethics committee

#### Experimental task and procedure

This was similar to Experiment 1 except that here, instead of reporting their belief of success, participants were instructed to report their decision to engage a focussed attention state in the upcoming trial (low vs high).

#### Machine learning protocols

These were similar to Experiment 1 except that here we employed the logistic regression to predict the decision to engage focussed attention (i.e. low vs high).

### Results

#### Classification results

First, we report the pairwise correlations between the features used for classification: (i) awareness and confidence: 0.25873 +/- 0.255267, t(15) = 3.81, p = 0.00515; (ii) correctness and awareness: 0.267667 +/- 0.116857, t(15) = 8.23, p = 0.00000182; (iii) correctness and confidence is 0.20129 +/- 0.157274, t(15) = 4.88, p = 0.00059. There is room therefore for each variable to provide distinctive information for the classification of the decision to engage attention.

We then employed a logistic regression classifier to predict the decision to engage attention on a given trial based on awareness, confidence, and correctness features from the preceding trials. We note that the probability of a decision to adopt a focussed attention state was 0.5005 +/- 0.0510, which is not different from 0.5 (p = 0.86) and hence indicates that the likelihood of each decision to engage attention was similar. We found that the decision to engage a particular state of attention could be significantly predicted above chance levels using information from the previous trials. As shown in Figure 5, classification accuracy was highest when information from the previous trial was used. Prediction accuracy dropping close to chance level based on information from 3 trials back. P-values following the permutation analyses for the different time windows were as follow: 1-back: p < 0.0025, 2-back: p < 0.0081, 3-back: p = 0.268, 4-back: p < 0.022. Similar results were obtained with the Random Forest classifier (see Supplemental Figure 5).

**Figure 5:**
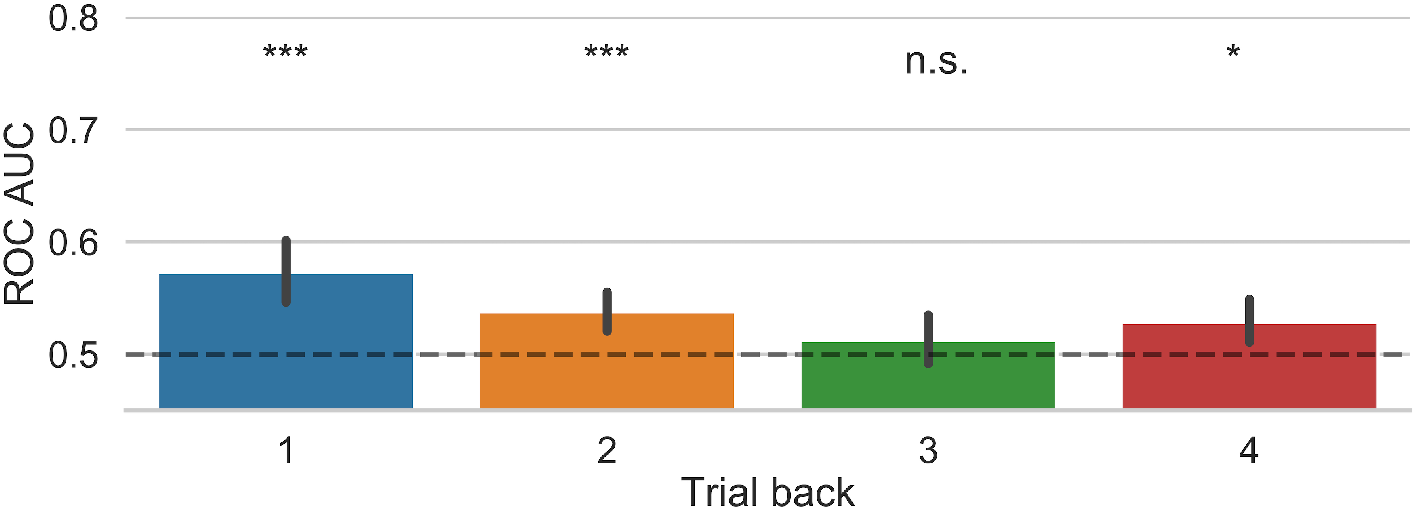
Results from the logistic regression classification model of the decision to engage attention tested separately for each of trials back. A value of 1 indicates that the odds of predicting the class is the same regardless of the feature value. Error bars represent bootstrapped 95% confidence intervals.n.s (not significant); * p < 0.05, *** p < 0.001.

Next, we assessed the relevance of each of the different attributes (awareness, confidence, and correctness) for the classification. Similar to Experiment 1, we performed an ANOVA over the odd ratios of the logistic regression with time window (1,2, 3 and 4 trials back) and attributes as factors. The results showed a main effect of time window, F(3,45) = 3.224, p = 0.0313, *η*^2^ = 0.043, but there was no main effect of attributes, F(2,30) = 2.052, p = 0.146, *η*^2^ = 0.023, and no interaction, F(4,60) = 0.97, p = 0.45, *η*^2^ = 0.023. However, Figure 6 shows that at least when observers rated high confidence on the previous trial, the probability of a decision to engage a focussed attention state on the next trial increased by 1.45, which was significantly different from correctness (p = 0.03, corrected). A similar result was obtained with the Random Forest classifier; however this also showed that awareness was more important for the classification than correctness (see Supplemental Figure 6).

**Figure 6:**
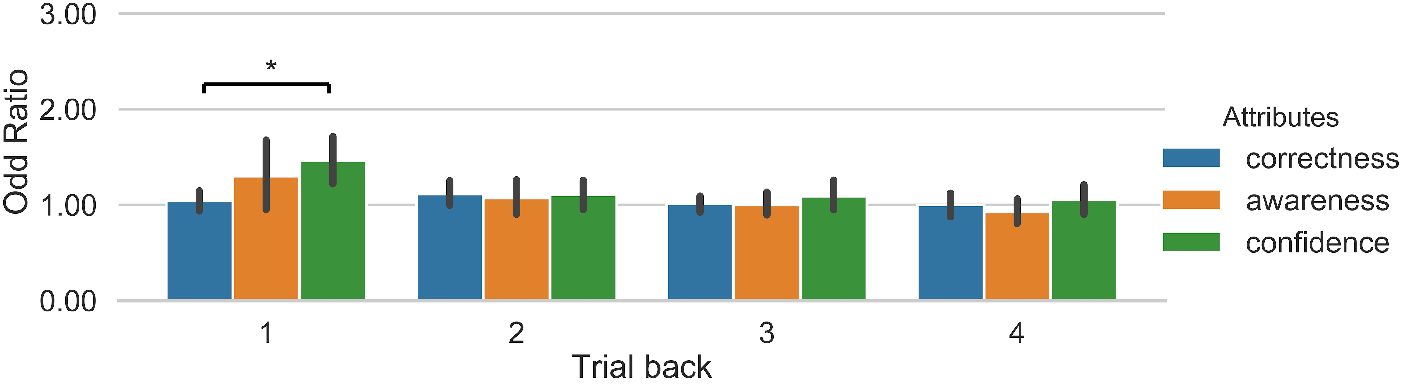
Illustration of odd ratios from the logistic regression for the different features involved in the classification of the decision to engage attention. Odd ratios were computed separately for each of trials back. Error bars represent bootstrapped 95% confidence intervals. * p < 0.05.

#### Univariate results

We also performed univariate analyses of the probability of a decision to engage a focussed attention state a as function of each level of correctness, awareness and confidence on the previous trial. Because the multivariate classification results were strongest based on information from 1 trial back, we conducted the univariate analyses for 1 trial back to assess the commonalities between the approaches. Accordingly, we conducted a 2 (Correctness: Hit, Error) x 2 (Awareness: Aware, Unaware x 2 Confidence (high, low) repeated measures ANOVA on the probability ofa decision to engage focussed attention. There was no main effect of correctness on the probability of a belief of high success (F(1,13) = 0.25, p = 0.625), no main effect of awareness (F(1,13) = 0.607, p = 0.45) and no main effect of confidence (F(1,13) = 4.463, p = 0.055). There were no interactions between factors F(1,13) = 0.901, p = 0.360, for awareness and correctness; F(1,13) = 0.133, p = 0.721, for correctness and confidence; F(1,13) = 0.994, p = 0.337, for awareness and confidence, and F(1,13) = 0.371, p = 0.553, for the three-way interaction. The probability of a decision to engage attention in the next trial did not appear to be affected by the visual awareness state on the previous trial, not by the actual accuracy in the task. Only the main effect of confidence marginally approached significance; higher confidence in the previous trial was associated with decisions to invest more attention in the next trial. This is consistent with the classification analyses reported above, however it is clear that the multivariate classifier in the context of cross-validation was able to capture more information than univariate analyses based on significance testing of probability difference in which all data is used at once to fit the model.

Further, we performed a linear mixed regression analysis to predict the prospective decision to engage attention with fixed effects for each of the confidence, awareness and accuracy attributes considering the recent trial history up to 4 trials back with lagged factors and random intercepts for each participant. The results showed that confidence ratings from 1-trial back predicted the decision to engage attention in the current trial (see Supplemental Figure 7). While the classification results showed that information up to 2-trials back predicted the decision to engage attention (Figure 5), the present results are nevertheless in keeping with the analysis of the odd ratios of the logistic regression classifier which also indicated that confidence of 1-trial back significantly predicted the current decision (Figure 6).

#### Signal detection results

A 2 by 2 ANOVA with awareness (unaware, aware) and decision to engage a focussed state of attention (low, high) as factors was carried out on the observer’s perceptual sensitivity to the orientation of the Gabor. There was an effect of awareness on type-1 d’ (F(1,15) = 69.53, p < 0.000000515, *η*^2^ = 0.462682189). Perceptual sensitivity was higher on aware relative to unaware trials. There was no effect of the decision to engage attention on type-1 d’ (F(1,15) = 1.388, p = 0.257, *η*^2^ = 0.0054) and no interaction effect between decision to engage and awareness (F(1,15) = 1.877, 0.191, *η*^2^ = 0.00216).

A similar ANOVA was carried out on metacognitive sensitivity scores. There was an effect of awareness on meta-d’ (F(1,15) = 7.084, p = 0.0178, *η*^2^ = 0.065). Metacognitive sensitivity was slightly better on aware relative to unaware trials, though it remained significantly above chance in the unaware trials (i.e. t(15) = 2.88, p < 0.011 and t(15) = 3.47, p < 0.003, for the decisions to engage less and more attention respectively). There was also an effect of the decision of engage attention on meta-d’ (F(1,15) = 8.824, p = 0.00953, *η*^2^ = 0.0631). Metacognitive sensitivity was better following decisions to engage a more focussed attention state. There was also an interaction between the factors (F(1,15) = 5.279, p = 0.0364, *η*^2^ = 0.027). Pairwise comparisons showed that on aware trials, meta-d’ was better when participants decided to adopt a more focussed attention state (t(15) = 2.79, p < 0.014). This was not the case when the participants reported being unaware of the target (t(15) = 1.772, p = 0.097). These results are depicted in Figure 7. This result indicates that the decision to attend had an effect on metacognitive sensitivity. We note, however, there were no effects of the decision to engage attention on M-ratio (F(1,15) =2.55, p=0.13). There was also no effect of awareness (F(1,15) = 0.005, p = 0.94) and no interaction between the decision to engage and the state of visual awareness on M-ratio (F(2,15)=0.502, p=0.48). However, we believe it is unlikely that the influence of the decision to engage attention in meta-d’ can be explained by differences in d’. We shall come back to this point in the Discussion.

**Figure 7:**
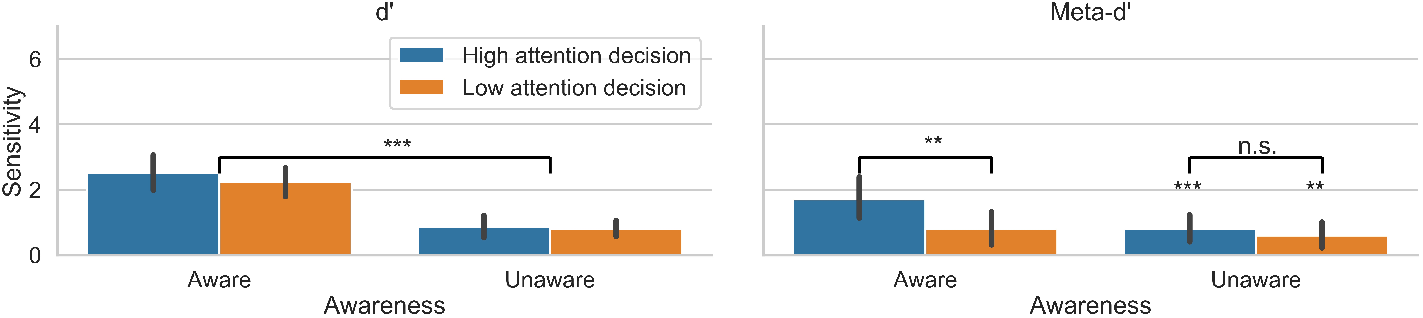
Perceptual sensitivity and metacogntive sensitivity scores as function of the state of awareness and the decision to engage attention. Error bars represent standard error of the mean. * p < 0.05, ** p < 0.01, *** p < 0.001.

## Discussion

The results showed that decisions to engage a focussed attentional state do not seem to affect perceptual sensitivity, at least in the current experimental conditions with single targets and no competing distracters. However, decisions to engage in a more focussed attention state influenced the observer’s metacognitive sensitivity. Although there was no effect of the decision to engage attention on M-ratio, we think it is unlikely that the effect of the decision to engage in meta-d’ can be accounted for in terms of differences in d’ given that these were only minimal and statistically insignificant. Most critically, previous research indicates that d’ and meta-d’ can be sensitive to different factors. For instance, using a similar paradigm to the one used in this study, we have previously shown that visibility has a strong effect on d’ but not on meta-d’,^11^ indicating that meta-d’ and d’ are not necessarily based on a similar process, or operate on a similar type of information (see also^22^).

This result is also in keeping with evicendece that that metacognitive confidence and task performance are dissociable.^22, 29–31^ Hence, we think the application of M-ratio in the current experimental context may not be optimal. In any case, the data indicate that the decision to engage attention had an effect on meta-d’, while this was not the case for prospective beliefs of success in Experiment 1. This suggests that decisions to engage attention may influence the observer’s evaluation of the correctness of perceptual decisions, although we note that no effects were seen in M-ratio.

On the other hand, the present results replicate the findings of Experiment 1 concerning the prediction of the future prospective belief of success, but crucially in a new decision context related to the attention state for the upcoming task. Intriguingly, we observed that high confidence in the previous trial predicted decisions to engage a more focussed attentional state in the next trial. Yet the opposite might be argued, namely, that when observers have held low confidence or have been incorrect, they would then decide to engage more focussed attention on the next trial. This is because errors or low confidence may in principle prompt subjects to be more cautious and try to boost attention to improve performance in the next trial. However, this is not always the case^32^ and it has been argued that it may depend on the type of error.^33, 34^ A related issue is the extent to which decisions to attend are metacognitive in nature. We think that prospective decisions are metacognitive insofar observers monitor their own recent behavioural performance and attentional state in order to engage in a cognitive setting that can promote more adaptive behaviour in the next trials. However, we acknowledge that our study is not well suited to address this issue given that we did not collect information from our participants regarding the type of strategies they used to make prospective decisions to attend.

We now turn to integrate the findings from the two experiments.

### Across-experiment Generalization Results

Finally, we analyzed the data from both experiments together in order to estimate how much information learned from one experiment can be transferred to the other experiment. We fitted the models with the features in one experiment (i.e. prospective of belief of success), and then test the trained model in the other experiment. To estimate the variance of the cross-experiment generalization, we conducted a cross-validation procedure as follows. First, we selected one of the experiment as the source and the other experiment as the target. Second, we preprocessed both the source and target experiment as described in the preprocessing section. Data from all the subjects were concatenated as one whole dataset. Third, the cross-validation method described above was applied to both the source data and the target data, while the training data was sampled from the source data and the testing data was sampled from the target data. Particularly, in each fold, 80% of the source data formed the training set and 20% of the target data formed the testing set. With 100 iterations of the cross-validation procedure, we estimated the variances of the transfer learning given N-back trials (N = 1, 2, 3, 4).

We first trained the classifiers using the data from the prospective belief of success (Experiment 1) and tested the classifiers using the data from the prospective decision to engage attention (Experiment 2). We found that the logistic regression model was able to decode the decision to attend based on the pattern of awareness, confidence, and correctness in the previous trial of the belief of success experiment (p = 0.0106), but not with the attributes in 2, 3, or 4 trial back (p > 0.9).

We then trained the classifier using the data from the prospective decision to engage attention (Experiment 2) and tested the classifier using the data from the prospective belief of success (Experiment 1). The classifier was able to decode the prospective belief of success based on the pattern of awareness, confidence, and correctness in the previous trial of the decision to attend experiment (p = 0.0004), but this was not the case for N = 2 or 3 or 4 trials back (lowest p-value = 0.0739).

These results are depicted in Figure 8. This pattern of results found with the random forest classifier was similar, though the random forest classifier trained with data from 3 trials back in Experiment 2 generalised to predict beliefs of success in the current trial of Experiment 1 (see Supplemental Figure 8).

**Figure 8:**
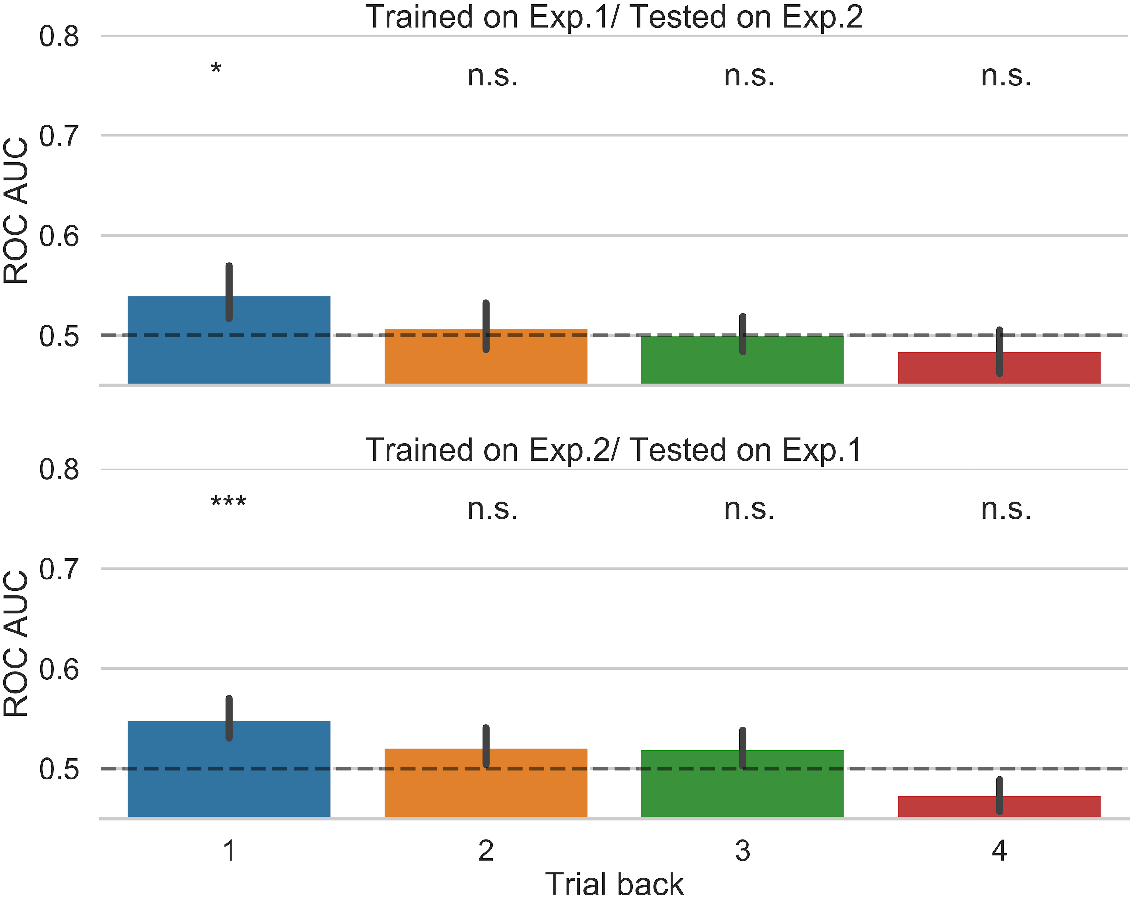
Across-experiment generalization results. Classifiers were trained on data from either experiment and tested on the other experiment. This was done separately for each of trials back. Error bars represent bootstrapped 95% confidence intervals. n.s (not significant); * p < 0.05, ** p < 0.01, *** p < 0.001.

We also performed a cross-experiment comparison on the effect of the prospective decision on metacognitive sensitivity. Recall that in Experiment 1 we did not find that the prospective belief of success had any influence on type-1 perceptual sensitivity or type-2 metacognitive performance. To quantify whether the result of Experiment 2 was significantly different from Experiment 1 we compared the pattern of metacognitive sensitivity across experiments. In particular, we used the aware trials - in which the effect of the decision to engage attention was found-to compute the difference in meta-d’ following decisions to adopt a more focussed relative to a less focussed attention state. The same was done in Experiment 1 for the high vs low prospective belief of success. An unpaired t-test showed that the influence of the decision to attend on meta-d’ was higher than the influence of the prospective belief of success (t(28)=2.06, p=0.048).

## General Discussion

We sought to understand the sources of information determining prospective decision making in the context of a perceptual task using standard machine learning techniques. Across two experiments, we found that a logistic regression classifier significantly predicted the upcoming prospective belief of success based on the pattern of awareness, confidence, and correctness exhibited in previous trials. Information from the previous trial led to the highest accuracy in the prospective belief of success, with classification accuracy increasingly dropping with up to four trials back. Both types of prospective decisions were predicted by the prior states of response confidence and visual awareness. Across the two experiments, we observed that task correctness was less important than both awareness and confidence for the classification of both prospective decisions. Importantly, the novel question addressed in the current study related to the cross-experiment generalization of the different prospective decisions. Accordingly, multivariate classification analyses were conducted for each trial back separately because this allowed us to assess whether a classifier trained in the context of one type of prospective decision (e.g. prospective confidence), in a particular trial back, generalised to predict a different type of prospective decision (i.e. to engage attention). We found that a classifier trained with information from 1 trial back was predictive of the other prospective decision, crucially, in a different set of participants which had only performed one of the experiments. There was no such generalization for 2, 3 and 4 trials back. We believe the absence of cross-experiment generalisation based 2, 3 or 4 trials likely reflects that just preceding decisions are most powerful to guide current decisions. This is in keeping with the study by Fleming and colleagues (2016) in which retrospective confidence judgments were more strongly related to information from the prior trial and a similar pattern was observed for prospective confidence. Strong lag-1 recency biases have also been reported in different experimental contexts beyond decision making. For instance, serial dependency effects in perceptual experience are strongest based on the similarity between the current stimulus and the stimulus of the previous trial.^35^ We also note that strong lag-1 dependency has also been observed in other dependent variables that are critical for confidence generation, such as the latencies of perceptual choices^36^ and also the latencies of confidence judgments.^37^ Hence we believe that a simple recency-bias is likely to explain why the predictions from the just the preceding trial are strongest.

The classification results conducted separately in each experiment with beliefs of success or decisions to engage showed that information from up to several trials back was predictive of the current prospective decision. However, the extent to which information from 3 or 4 trials back played a role in guiding the current prospective decision remains an outstanding question. These results merely show that there is information from up to 3 and 4 trials back that is associated with the current prospective decision. Yet, prediction scores and also the coefficients from linear models in which all data are fit at once, only provide information that is necessary but not sufficient to conclude that information from several trials back is genuinely causally involved in the formation of the current prospective decision, namely, that participants employ that past information to guide the current choice. This limitation may be addressed by devising novel paradigms to experimentally manipulate confidence and awareness and assess the effects on subsequent prospective decisions. Additional work is needed to make further determinations.

Previous research has shown that components of retrospective confidence estimates (e.g. mean, variance) are highly correlated across testing sessions involving a similar experimental task, although the generalization of confidence components across different task contexts was weaker.^38^ Additional studies support the view that the precision of retrospective metacognitive judgments correlates across perceptual and memory tasks^39, 40^ and across sensory modalities.^41^ Other studies have shown that observers use a similar confidence scale for different tasks of the same or different modality^42, 43^ and that confidence estimates on a given trial of a task carry-over to subsequent trial of a different task.^44^ Across the two experiments of the present study, information related to visual awareness and response confidence on the most recent trials determines future prospective decision making, regardless of the type of prospective judgment (i.e. prospective belief of success or a decision to engage attention). The results indicates that a common pattern of information underlies the formation of seemingly different prospective decisions. This conclusion is also supported by cross-experiment generalization results.

However, despite the common information pattern based on past confidence and awareness that underlies the formation of prospective judgments, only the prospective decisions to attend appeared to influence type-2 performance, namely, the observer’s retrospective evaluation of the correctness of perceptual decisions (meta-d’), but this was not the case following a prospective belief of high success. From the perspective of the ‘self-fulfilling prophecy’, prospective beliefs of success may set an expectation concerning upcoming behavioural performance that the participant is motivated to meet^13, 14^ and accordingly, a belief of performance success might in principle encourage observers to invest more cognitive resources in the upcoming trial. Our results suggest that this is not the case.

Prospective beliefs of success concerned here a low-level perceptual discrimination task. It is possible that prospective beliefs are more diagnostic of the upcoming behavioural performance in different task domains, namely, memory.^4, 45^ Another possibility is that decisions to engage attention are more likely to be embodied by comparison to beliefs of success. Accordingly, recent theoretical frameworks borrowing from ecological psychology^46^ propose that perceptual biases and decisions are not independent of action. In this framework, perception drives decisions and action, but actions also drive subsequent experiences in a dynamic perception-action loop.^47^ We propose that decisions to engage with the environment (i.e. to deploy a focussed attention state) are more likely to be embodied in the action system and hence are very likely to trigger commitment towards that action, while prospective beliefs may not. It is possible that decisions to engage attention trigger preparatory control which in turn can influence subsequent cognitive processing. Decisions to engage attention influenced participants evaluation of the correctness of perceptual decisions but had no effect on perceptual processing itself, however. The latter is likely due to the absence of visual competition in the displays with single targets at central fixation. It is well known that attention effects on visual processing are stronger when there is space- or object-based competition for selection.^48^

A important question is whether prospective decisions to engage attention are causally, rather than merely associated, with subsequent changes in behavioural performance. Our study did not manipulate confidence or awareness in a way that allowed us to infer causal relationships with subsequent behavioural performance. Hence the correlational nature of our design is a limitation of the present study which should be addressed in future work. However, there are grounds to argue that prospective decisions can play a causal role in behavioural control. First, this view is consistent with the theoretical framework outlined above that proposes the existence of dynamic perception-decision-action loops. Accordingly, causal associations between prospective decisions and subsequent cognitive and action control processes are expected within this framework. Moreover, in the memory domain, judgments of learning are typically thought to play an important role in how individuals control their own learning; similar effects, even stronger, have been recently reported for retrospective confidence judgements.^49^ More relevant for the present study is recent work showing that prospective confidence estimates are associated with neural markers of proactive control.^50^ In this study, participants performed a color judgment task in a multi-item display followed by a confidence rating. This task was preceded by visual cues that were paired with target displays, which in the critical medium difficulty conditions (i.e. colour search displays with low mean-low variance and high mean-high variance) were associated with different levels of confidence during task performance, while crucially the level of performance accuracy was matched. On a different type of trials, participants were only presented with the cues and then rated the confidence associated with performing the task, as if the stimulus was presented. The results showed that participants learned the confidence associated with each cue, and this estimate was similar to the confidence rated following task performance. Notably, EEG markers of preparatory control and evidence accumulation were influenced by the level of predicted confidence;^50^ however, little evidence for this was found in the critical medium difficulty trials, namely, EEG markers of preparatory control were influenced by the level of predicted confidence in the most easy and hard conditions, which were also associated with performance differences. Hence, while there is some suggestive evidence that prospective decisions may be causally involved in the control of behaviour, our study only found weak evidence for this (i.e. the prospective decision to engage attention was associated with better type-2 metacognitive performance but there were no effects on type-1 performance).

In summary, the present study indicates that a common representational structure supports the dynamic formation of seemingly different types of prospective judgements. Additional research is however needed to test whether these observations generalize to different task contexts and cognitive domains beyond perceptual decision making.

## Supporting information

Supplementary Results

## Acknowledgements

D.S. acknowledges support from the Basque Government through the BERC 2018-2021 program, from the Spanish Ministry of Economy and Competitiveness, through the ‘Severo Ochoa’ Programme for Centres/Units of Excellence in R & D (SEV-2015-490) and also from project grants PSI2016-76443-P from MINECO and PI-2017-25 from the Basque Government.

## Data availability statement

Data and code are available at https://github.com/nmningmei/metacognition

## Author contributions

N.M. performed data analyses and wrote the paper; S.R. and E.O. run the experiments and contributed towards data analyses; D.S. designed the study, wrote the paper, and supervised all aspects of the project.

## Competing interests statement

The authors declare no competing interests.

