## Supplementary Results for "Similar history biases for distinct prospective decisions of self-performance"

**Random forest classification results:** A regularized random forest (Scikit-learn implementation). The implementation utilized the cross-entropy as an objective function with 500 decision trees. This number of decision trees was used as part of the regularization in order to avoid overfitting and also to approximate a simpler model which would enable comparison with the results from the logistic regression. Tree-based models are composed of nodes with each representing a level of depth in the model resulting from binary decision nodes where each node compares one feature value of the samples to a threshold. The maximum depth of each decision tree classifier was set to 1 so that only 1 best feature would be eventually used to make the decision. This means that during the training, with bootstrapped samples and features, the individual decision tree classifier ranks the importance of the feature by dropping one of the features and estimate a new decoding score after the dropping. The more loss in the decoding score compared to the original score, the more important the given feature was. In other words, we aimed to estimate the best feature among all features used (i.g confidence, awareness, and correctness). The majority rule was then applied to estimate the feature importance across all decision tree classifiers.

Note the sum of feature importance of all features add up to 1 in the implementation of Scikit-learn: (https://scikit-learn.org/stable/modules/generated/sklearn.ensemble.RandomForestClassifier.html#sklearn. ensemble.RandomForestClassifier.feature_importances_), and so this will be the case across all trials back. Hence, the main effect of the time window in the ANOVA is not meaningful because the average feature importance at each time window would be exactly the same (i.e. 0.33) and consequential it is not reported.

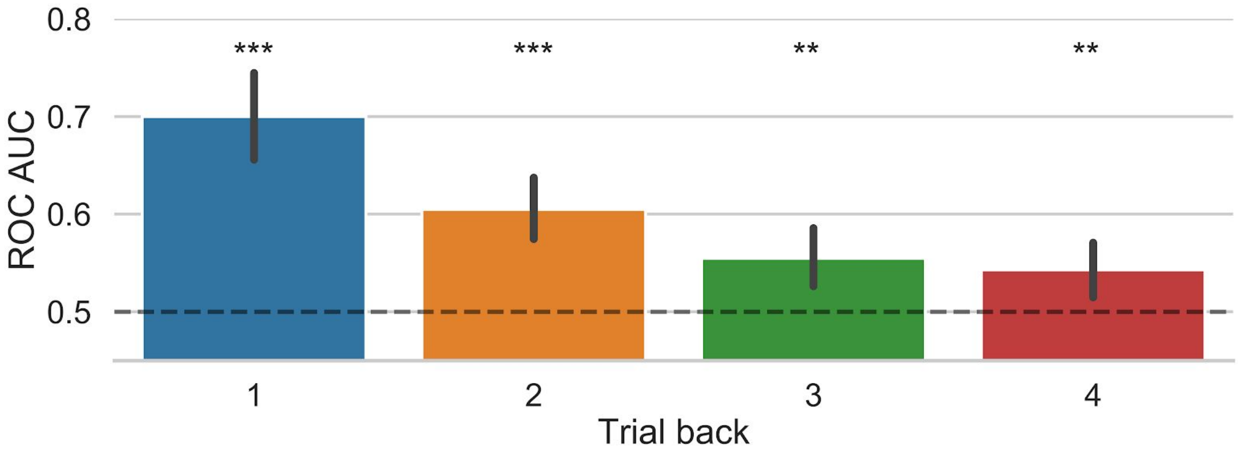


**Supplemental Figure 1.** The prediction of success could be classified with the highest accuracy by using the features from the previous trial using the Random Forest Classifier, with prediction accuracy dropping close to the chance level based on information from 4 trials back. P-values following the permutation test for the different classification analyses were: 1-trial back: p <

0.0004, 2-back: $p < 0.0004$, 3-back: $p < 0.0036$, 4-back: $p < 0.0076$. Error bars represent bootstrapped 95% confidence intervals, resampled from the distribution of decoding scores of individual participants with 1000 iterations. ** $p < 0.01$, *** $p < 0.001$.

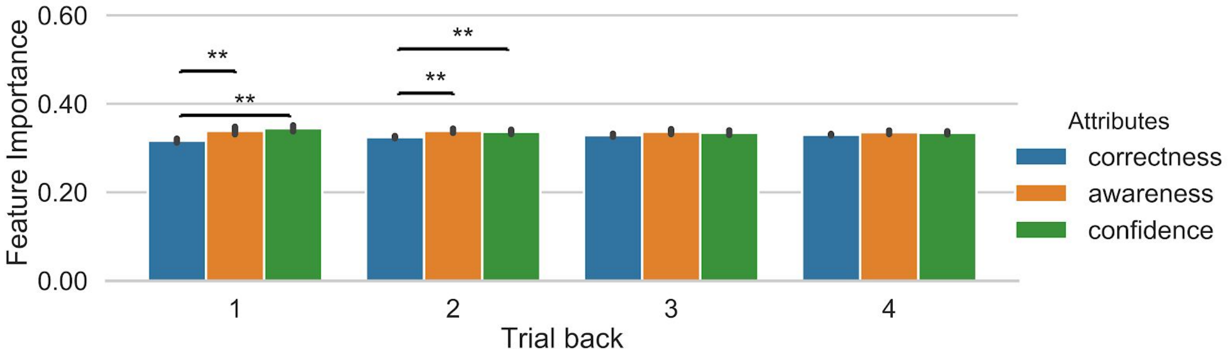


**Supplemental Figure 2**. The analysis of feature importance showed a significant main effect of attributes, $F(2,28) = 8.244$, $p = 0.00513$, $\eta2 = 0.232$). Post-hoc permutation paired t-tests showed that both confidence and awareness ($p = 0.0003$ and $p = 0.0023$) have a higher feature importance relative to correctness. There was also a significant interaction between factors, $F(6,84) = 6.063$, $p = 0.000263$, $\eta^2 = 0.112$), showing that the relative importance of confidence and awareness vs. correctness for the classification decreased with the number of previous trials. Post-hoc t-tests were conducted to further assess this interaction. For 1-trial back, confidence ($p = 0.0012$) and awareness ($p = 0.0020$) had different feature importance than correct, but there was no difference between confidence and awareness ($p = 1.0$). For 2-trials back, confidence ($p = 0.0063$) and awareness ($p = 0.0020$) had different feature importance than correctness, but there was no difference between confidence and awareness ($p = 1.0$). This pattern of results was not observed for 3 and 4-trials back (all ps > 0.2761). Error bars represent bootstrapped 95 % confidence intervals. ** $p < 0.01$.

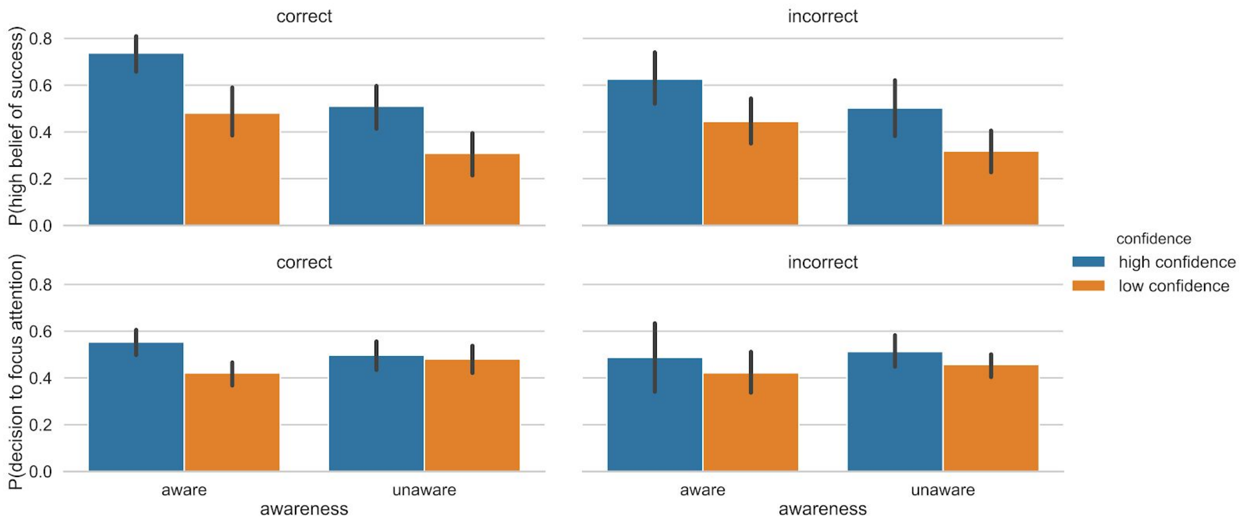


**Supplemental Figure 3**. (upper panel) Illustration of the probability of high belief of success and (lower panel) probability of a decision to engage focussed attention, as a function of the correctness, awareness, and confidence on the previous trial. Error bars represent bootstrapped 95% confidence intervals.

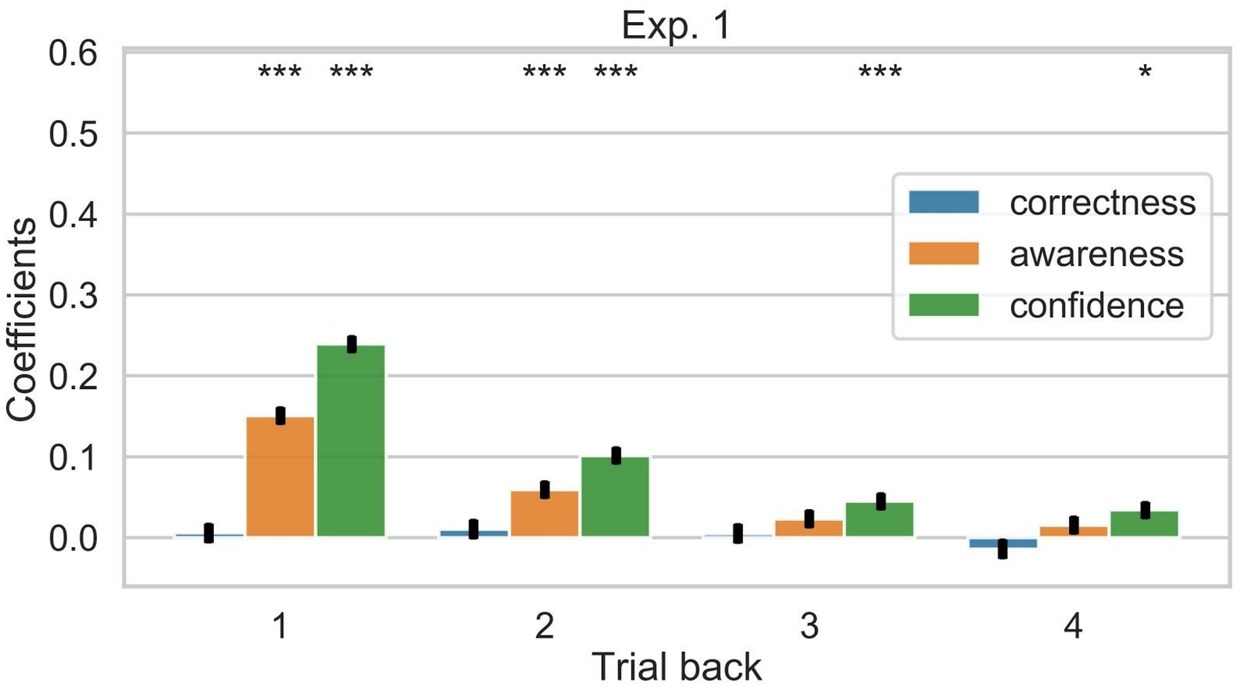


**Supplemental Figure 4.** A mixed-linear regression analysis was conducted to predict the prospective belief of success with fixed effects for each of the confidence, awareness and accuracy attributes considering the recent trial history up to 4 trials back for each attribute (12 regressors in total) and random intercepts for each participant. Error bars represent the standard error of the estimated coefficients. One-sample t-tests against zero assess the statistical significance of each coefficient. The results reported here are corrected for multiple tests using Bonferroni. The coefficients of 1-trial back were significant for confidence ($p = 1.352e-110$) and awareness ($p = 1.429e-42$) and the same pattern was observed for the coefficients of 2-trials back ($p = 6.918e-21$ and $p = 5.784e-07$). The coefficient of confidence were reliable at 3-trials back ($p = 2.412e-04$) and also at 4-trials back ($p = 1.409e-02$); * $p < 0.05$, *** $p < 0.001$.

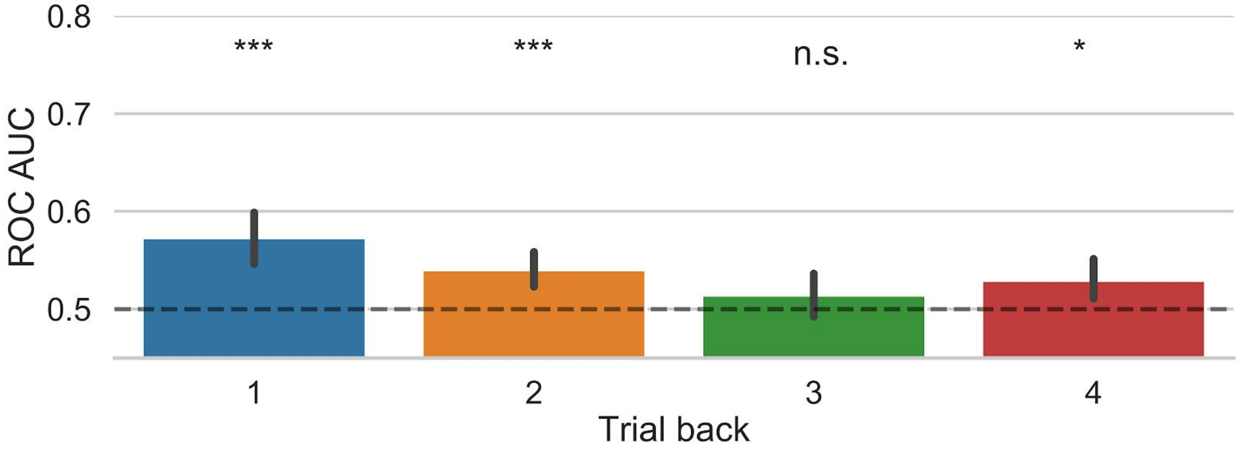


**Supplemental Figure 5**. We employed the random forest classifier to predict the decision to engage attention based on awareness, confidence, and correctness feature from the preceding trials. Same as logistic regression reported, the particular decision to engage a focused state of attention could be significantly predicted above chance level using information from the previous trials: 1-trial back: $p < 0.0004$, 2-back: $p < 0.0005$, 3-back: $p = 0.3959$, 4-back: $p = 0.0138$ Error bars represent bootstrapped 95 % confidence intervals. n.s (not significant); * $p < 0.05$, *** $p < 0.001$.

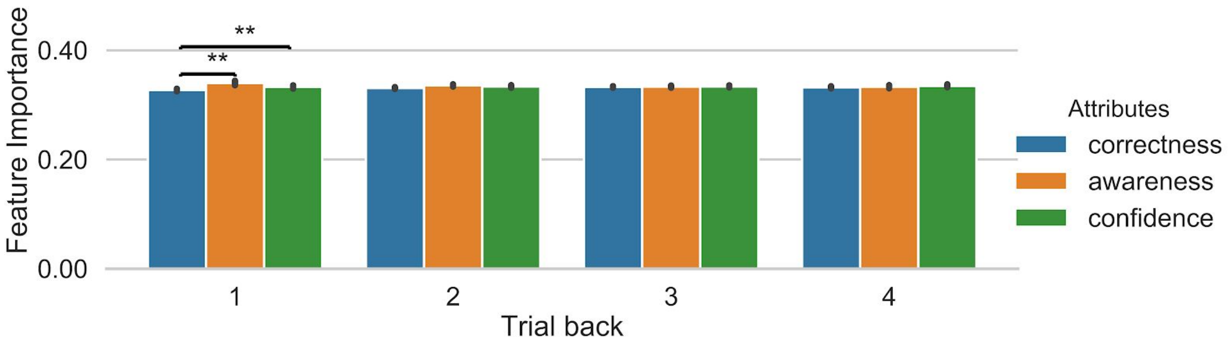


**Supplemental Figure 6**. The analysis of feature importance showed a significant main effect of attributes, $F(2,30) = 5.378$, $p = 0.0101$, $\eta 2 = 0.103$). Post-hoc permutation paired t-tests showed that both confidence and awareness ($p = 0.0061$ and $p = 0.0075$) have a higher feature importance relative to correctness. There was also a significant interaction between time window and attributes, $F(6,90) = 3.667$, $p = 0.00265$, $\eta^2 = 0.119$, showing that the relative importance of confidence and awareness vs. correctness for the classification decreased with the number of previous trials. Post-hoc t-tests were conducted to further assess this interaction. For 1-trial back, confidence ($p = 0.0034$) and awareness ($p = 0.0047$) had different feature importance than correctness, but there was no difference between confidence and awareness ($p = 1.0$). This pattern of results was not observed for 2, 3 and 4 trials back (all ps > 0.0729). Error bars represent bootstrapped 95 % confidence intervals. ** $p < 0.01$.

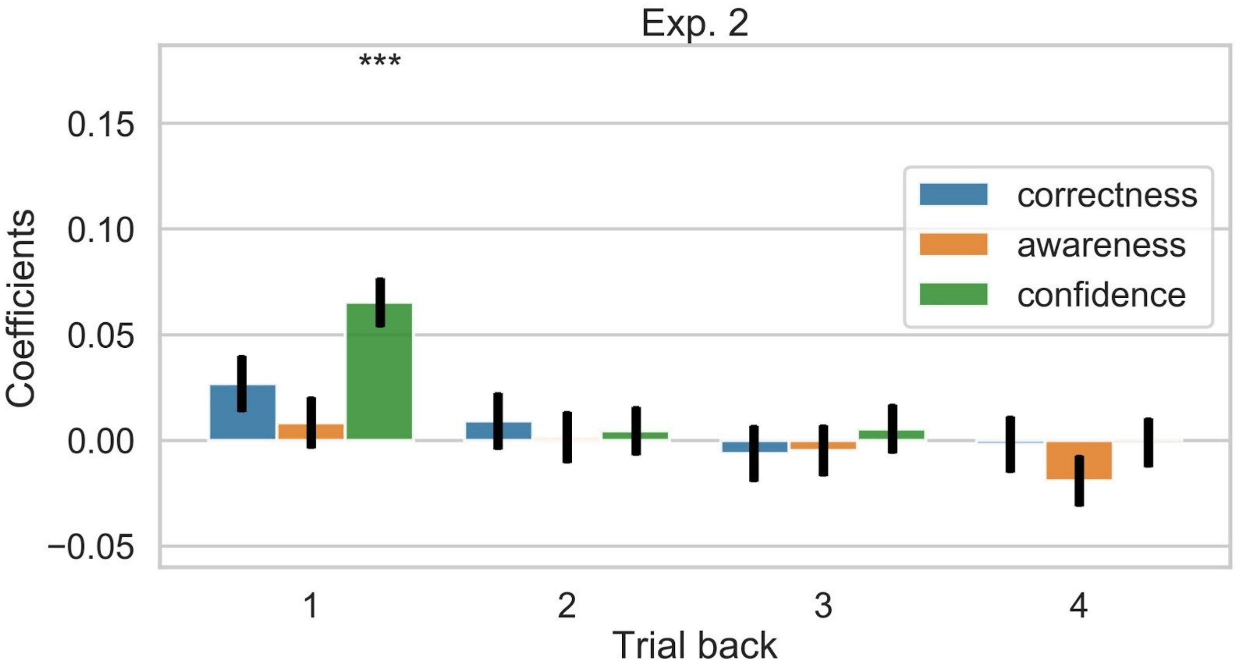


**Supplemental Figure 7**. A mixed regression analysis was conducted to predict the decision to engage attention with fixed effects for each of the confidence, awareness and accuracy attributes considering the recent trial history up to 4 trials back for each attribute (12 regressors in total) and random intercepts for each participant. Error bars represent the standard error of the estimated coefficients. One-sample t-tests against zero assess the statistical significance of each coefficient. Only the coefficient of confidence of 1-trial back was significant ($p = 1.781e-07$). Results are corrected for multiple tests using Bonferroni; *** $p < 0.001$.

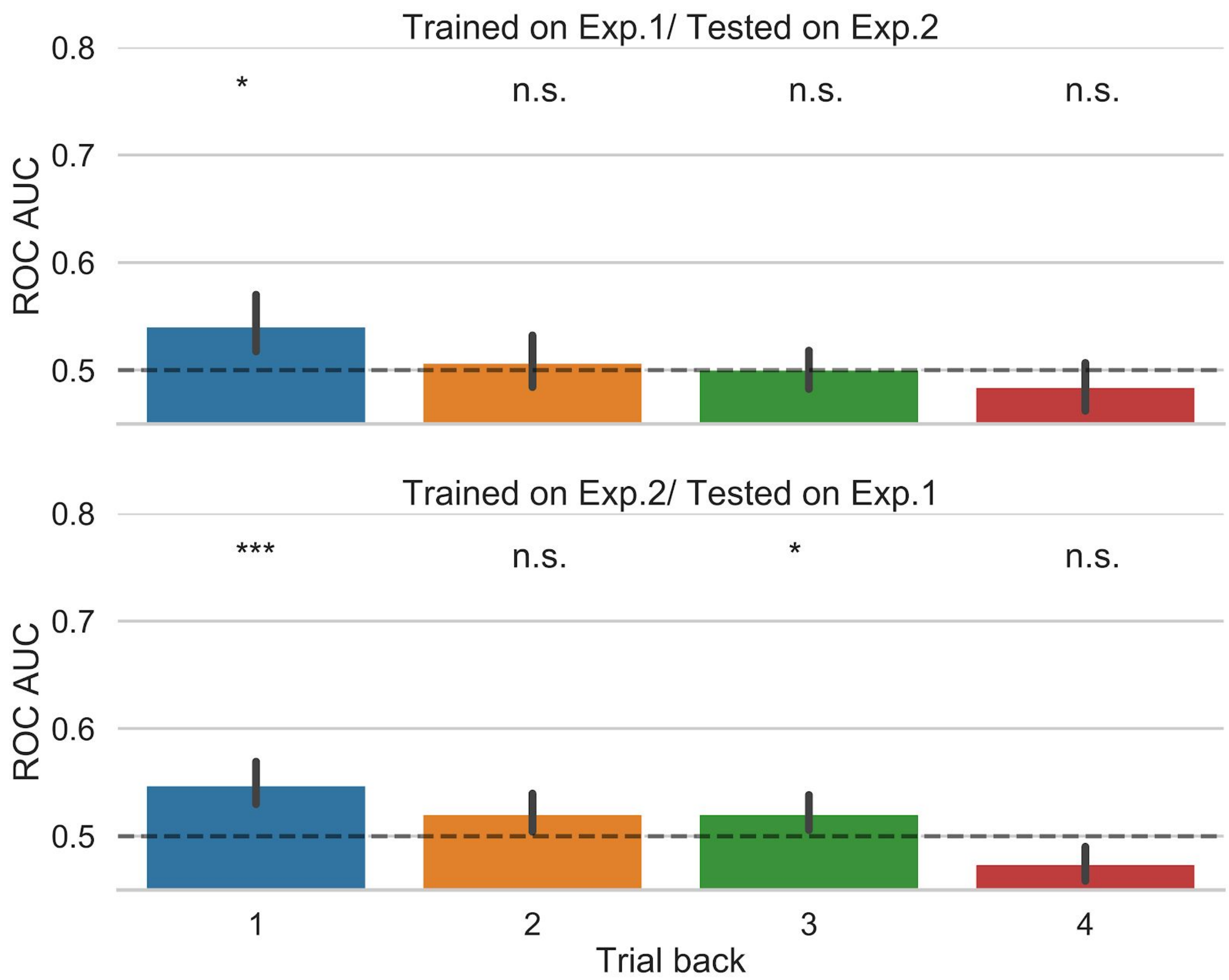


**Supplemental Figure 8**. Decoding results of the random forest classifier cross-generalizes between 2 experiments. When trained on experiment 1 and tested on experiment 2, only decoding results of the 1-trial back is significantly higher than chance level, $p = 0.0105$. When trained on experiment 2 and tested on experiment 1, decoding results of 1-trial back ($p = 0.0005$) and 3-trial back ($p = 0.0369$) trial are significantly higher than chance. Error bars represent bootstrapped 95 % confidence intervals. n.s (not significant); * $p < 0.05$, *** $p < 0.001$.
